# The neural features in the precentral gyrus predict the severity of internet game disorder: results from the multi-voxel pattern analyses

**DOI:** 10.1101/2020.08.26.267989

**Authors:** Shuer Ye, Min Wang, Qun Yang, Haohao Dong, Guang-Heng Dong

## Abstract

**Importance:** Finding the neural features that could predict internet gaming disorder severity is important in finding the targets for potential interventions using brain modulation methods.

**Objective:** To determine whether resting-state neural patterns can predict individual variations of internet gaming disorder by applying machine learning method and further investigate brain regions strongly related to IGD severity.

**Design:** The diagnostic study lasted from December 1, 2013, to November 20, 2019. The data were analyzed from December 31, 2019, to July 10, 2020.

**Setting:** The resting-state fMRI data were collected at East China Normal University, Shanghai.

**Participants:** A convenience sample consisting of 402 college students with diverse IGD severity

**Main Outcomes and Measures:** The neural patterns were represented by regional homogeneity (ReHo) and the amplitude of low-frequency fluctuation (ALFF). Predictive model performance was assessed by Pearson correlation coefficient and standard mean squared error between the predicted and true IGD severity. The correlations between IGD severity and topological features (i.e., degree centrality (DC), betweenness centrality (BC), and nodal efficiency (NE)) of consensus highly weighted regions in predictive models were examined.

**Results:** The final dataset consists of 402 college students (mean [SD] age, 21.43 [2.44] years; 239 [59.5%] male). The predictive models could significantly predict IGD severity (model based on ReHo: *r* = 0.11, *p*(r) = 0.030, SMSE = 3.73, *p*(SMSE) = 0.033; model based on ALFF: *r*=0.19, *p*(*r*) = 0.002, SMSE = 3.58, *p*(SMSE) = 0.002). The highly weighted brain regions that contributed to both predictive models were the right precentral gyrus and the left postcentral gyrus. Moreover, the topological properties of the right precentral gyrus were significantly correlated with IGD severity (DC: *r* = 0.16, *p* = 0.001; BC: *r* = 0.14, *p* = 0.005; NE: *r* = 0.15, *p* = 0.003) whereas no significant result was found for the left postcentral gyrus (DC: *r* = 0.02, *p* = 0.673; BC: *r* = 0.04, p = 0.432; NE: *r* = 0.02, *p* = 0.664).

**Conclusions and Relevance:** The machine learning models could significantly predict IGD severity from resting-state neural patterns at the individual level. The predictions of IGD severity deepen our understanding of the neural mechanism of IGD and have implications for clinical diagnosis of IGD. In addition, we propose precentral gyrus as a potential target for physiological treatment interventions for IGD.

**Key Points:** *Question:* Can machine learning algorithms predict internet gaming disorder (IGD) from resting-state neural patterns?

*Findings:* This diagnostic study collected resting-state fMRI data from 402 subjects with diverse IGD severity. We found that machine learning models based on resting-state neural patterns yielded significant predictions of IGD severity. In addition, the topological neural features of precentral gyrus, which is a consensus highly weighted region, is significantly correlated with IGD severity.

*Meaning:* The study found that IGD is a distinctive disorder and its dependence severity could be predicted by brain features. The precentral gyrus and its connection with other brain regions could be view as targets for potential IGD intervention, especially using brain modulation methods.

## Introduction

Internet gaming disorder (IGD) has been recognized as a serious mental health problem that is characterized by obsession with gaming, hypersensitivity to game cues and failure to resist the impulse to play games despite negative consequences ^1-3^. In recent decades, there has been a steady increase in IGD around the world, and an increased number of studies have focused on behavioral characteristics associated with IGD ^4-6^. In 2013, IGD was included in Section III of the DSM-5 (The Diagnostic and Statistical Manual of Mental Disorders, Fifth Edition) as a condition requiring future research ^7^. In 2015, gaming disorder was officially listed in the new version of the International Classification of Diseases (https://www.who.int/news-room/q-a-detail/gaming-disorder). Despite widespread interest in IGD, less is known about the neural substrates underlying this disorder.

The behavioral model of IGD has been well developed thus far. Extreme reward-seeking ^8^, defective executive inhibition ^9^, and risky decision-making ^10^ are commonly recognized as core components of IGD ^5, 11^. These findings provide a basis for clinical behavior therapy for IGD ^12, 13^. However, most of these cross-sectional studies revealed the static condition of IGD, which cannot explain the relationship between neural features and the degree of addiction. Thus, the exploration of neural substrates of IGD may advance our understanding of this mental disorder and provide new insights for diagnosis, intervention, and treatment.

Resting-state functional magnetic resonance imaging (rs-fMRI) is an effective task-independent method used to investigate the neural substrates of IGD. Two metrics of rs-fMRI were developed to measure neural activities in the human brain: (i) regional homogeneity (ReHo) measures regional synchronization at the whole-brain level and spontaneous local neural activity ^14^, and (ii) the amplitude of low-frequency fluctuation (ALFF) measures regional intensity of spontaneous fluctuations in the BOLD signal. These two neural indicators have been effectively used to explore the neural mechanism of IGD ^15-17^. For instance, ReHo alterations were found in regions related to sensory-motor coordination, audiovisual processing, and reward pathways in individuals with IGD compared with healthy controls ^18^. Additionally, studies that employed the ALFF method found abnormalities in the left medial orbitofrontal cortex, the left precuneus, the left supplementary motor area, the right parahippocampal gyrus, and the bilateral middle cingulate cortex among individuals with IGD ^19, 20^. Therefore, IGD has been widely proven to be related to abnormal brain activity, which leads to its addiction pattern ^21-23^. However, few studies have had a sample size large enough to ensure the reliability of their findings. More importantly, previous studies using conventional fMRI analyses were unable to effectively utilize massive fMRI data to delineate neural patterns and often focused on the differences between individuals with IGD and normal individuals at the group level.

Multivoxel pattern analysis (MVPA), a powerful data-driven machine learning method, is widely used in decoding brain activities and providing useful neural information to improve mental disorder diagnosis and treatment ^24, 25^. MVPA has unique advantages (e.g., having an increased sensitivity to detect brain patterns, allowing the extraction of feature weights, and characterizing neural code at the individual level) compared with conventional fMRI analyses. This method has been implemented to characterize neural coding and information processing in psychiatric studies ^26, 27^ and could be useful in identifying the neural features of IGD. However, to date, only one study has explored the neurobiological mechanism of IGD by using MVPA ^28^, suggesting that MVPA provides a potential way to distinguish individuals with IGD from recreational game users by decoding brain patterns represented by ReHo values. However, whether multiple brain activities can predict the IGD features of an individual remains unknown. Thus, determining whether different measures can predict IGD features and how they work is important and necessary.

Studies have demonstrated that addiction is related not only to abnormal brain activities in a specific region but also to atypical interactions between brain regions and networks ^22, 29^. Altered functional connectivity was shown between regions involving reward, cognitive processing, and executive control function in subjects with IGD ^30, 31^. Although MVPA based on local neural features can characterize brain patterns, it cannot be used to explore the regional relationships involved in IGD. Thus, graph theory and Granger causality analysis (GCA) were applied to address this problem. Graph theory provides a powerful framework to describe the whole brain topologically and has been widely used in studies about addiction ^32^. By constructing a network made up of nodes and edges, several topological metrics can be calculated to characterize regional properties and then reveal the potential significant position of the regions in the brain. Wang et al. ^33^ reported that subjects with IGD showed reduced node metrics in executive control and emotion-related regions, indicating the key roles these regions played in IGD. In addition, Granger causality analysis, which is a mathematical method used to build effective connectivity, can be applied to detect coupling among regions without assumptions about connections between them ^34, 35^. To date, two task fMRI studies have used GCA to explore the neural substrates of IGD. One study found that IGD severity was negatively correlated with connectivity from the middle frontal gyrus to the precuneus during a cue-carving task, and another study found abnormal effective connectivity within the salience network in adolescents with IGD ^36, 37^.

In the present study, we aimed to combine MVPA and graph theory analysis with GCA to decode the specific neural patterns of IGD in a large sample. We first applied the MVPA method to identify highly weighted regions as core brain areas in predicting IGD. The subsequent analysis revealed how these crucial regions for IGD work and interact with other regions. Specifically, we used MVPA to examine whether ReHo and ALFF could predict individuals’ IGD severity and to identify relatively highly weight brain regions that contributed to the model. Then, graph theory analysis was employed to further confirm the importance of selected regions in the IGD brain networks. Finally, GCA was implemented to explore how these regions interact with other brain regions in contributing to IGD. Based on previous studies, we hypothesized that (1) the predictive model based on ReHo and ALFF could significantly predict IGD severity; (2) regions related to reward processing, sensory-motor coordination, and executive control would be highly weighted in prediction models; (3) these highly weighted regions are crucial in the IGD brain network; and (4) IGD severity is related to effective connectivity between these regions and other regions in the whole brain.

## Methods

### Participants

Four hundred and two right-handed participants (239 males; 21.43 ± 2.44 years old) were recruited by advertisement from June 2013 to December 2019. All participants were free of any personal or family history of psychiatric disorders as assessed by an exhaustive structured psychiatric interview. Participants’ demographics and internet addiction test (IAT) scores are shown in Tab. 1. The study was conducted in accordance with the 1964 Helsinki Declaration and its later amendments and was approved by the Ethics Committee of Zhejiang Normal University. Written informed consent was obtained from all participants before participating in this research.

**Table 1.** Participants demographics and IAT scores (N=402).

|  | Male<br>(N=239) |  | Female<br>(N=163) |  |
| --- | --- | --- | --- | --- |
|  | Mean | SD | Mean | SD |
| Age (years) | 21.78 | 2.67 | 21.07 | 1.98 |
| Education (years) | 14.67 | 1.40 | 14.48 | 1.09 |
| Gaming history (years) | 3.80 | 0.56 | 3.61 | 0.74 |
| Gaming playing per weak(hours) | 6.94 | 3.00 | 7.08 | 3.59 |
| IAT score | 49.57 | 16.55 | 51.44 | 14.85 |

SD: standard deviation; IAT: Internet addiction test.

**Table 2.** Top 8 predictors for IGD severity and their relative weights in predictive power (percentage of the total weights) for predictive models based on ReHo and ALFF.

|  | Serial number <sup>a</sup> | Anatomical region <sup>b</sup> | MNI coordinates |  |  | Weight (%) | Size (voxels) | Hemi |
| --- | --- | --- | --- | --- | --- | --- | --- | --- |
|  |  |  | x | y | z |  |  |  |
| **ReHo model** | 52 | Postcentral gyrus | -54 | -9 | 23 | 1.67 | 19 | L |
|  | 75 | Postcentral gyrus | -38 | -27 | 60 | 1.57 | 19 | L |
|  | 35 | Precentral gyrus | 60 | 8 | 34 | 1.46 | 17 | R |
|  | 137 | Lateral occipital cortex | 45 | -72 | 29 | 1.25 | 19 | R |
|  | 77 | Postcentral gyrus | -24 | -30 | 64 | 1.17 | 19 | L |
|  | 62 | Precentral gyrus | -38 | -15 | 59 | 1.15 | 19 | L |
|  | 87 | Middle cingulate cortex | 8 | -40 | 50 | 1.12 | 19 | R |
|  | 132 | Precuneus | 11 | -68 | 42 | 1.12 | 19 | R |
| **ALFF Model** | 133 | Calcarine | 17 | -68 | 20 | 1.82 | 19 | R |
|  | 35 | Precentral gyrus | 60 | 8 | 34 | 1.63 | 17 | R |
|  | 147 | Calcarine | 9 | -76 | 14 | 1.56 | 19 | R |
|  | 52 | Postcentral gyrus | -54 | -9 | 23 | 1.44 | 19 | L |
|  | 148 | Cuneus | 15 | -77 | 32 | 1.35 | 19 | R |
|  | 86 | Postcentral gyrus | 34 | -39 | 65 | 1.33 | 19 | R |
|  | 67 | Supramarginal gyrus | -54 | -22 | 22 | 1.23 | 19 | L |
|  | 23 | Inferior frontal gyrus | -52 | 28 | 17 | 1.22 | 19 | L |

a: The serial numbers are taken from Dosenbach 160-node atlas.
b: The anatomical regions were defined based on Automated Anatomical Labeling (AAL) atlas.

### Measures

Participants were assessed for IGD severity with Young’s online Internet Addiction Test (IAT) ^38^. The IAT consists of 20 items related to internet use and internet-related addictive behavior. Each item can be rated on a 5-point Likert scale (from 1-rarely to 5-always). Higher IAT scores indicate greater internet use and addiction. The reliability and validity of the scale have been well validated ^39, 40^. In addition to the IAT, the DSM-5, which is a nine-item IGD diagnostic measure, was also applied to assess IGD severity ^2^. Participants were asked to answer “yes” or “no” to nine criteria occurring over the past 12 months. People who met more than 5 criteria were clinically diagnosed with IGD. Because the DSM-5 was published in 2014, only part of our participants (N = 365) took the assessment. IAT scores and DSM-5 scores are highly correlated.

### MRI data acquisition

Resting-state functional magnetic images were collected at East China Normal University using a 3T MRI system (Siemens Trio). The participants were simply instructed to keep their eyes closed and stay awake without performing any cognitive exercises. Head motion was minimized using foam padding and restraint. The imaging parameters were as follows: repetition time (TR) = 2000 ms, interleaved 33 slices, echo time (TE) = 30 ms, thickness = 3.0 mm, flip angle = 90°, field of view = 220 ×220 mm, and matrix = 64 × 64. Each fMRI scan lasted for 420 s and included 210 imaging volumes.

### Preprocessing

Preprocessing was conducted with DPABI v3.0 (Data Processing & Analysis for Brain Imaging: http://rfmri.org/dpabi), which is a pipeline toolbox for fMRI analysis in MATLAB ^41^. For each participant, the first 10 volumes were discarded to minimize the (transient signal) instability of the initial signal and adapt participants to the scanning environment (effect of scanner calibration). Subsequent data preprocessing included slice timing correction, head motion correction, spatial normalization to the standard MNI space with an EPI template and resampling into 3 × 3 × 3 mm^3^ voxels. The data used in the present study met the criteria of head motion <2.5 mm or 2.5°. Nuisance signals, including 24 motion vectors (i.e., six 6 head motion parameters, 6 head motion parameters one time point before, and the 12 corresponding squared items), the white matter signal, and the cerebrospinal fluid signal, were regressed out ^42^. Subsequently, the linear trends of time courses were removed, and the resulting images were temporally filtered with a bandpass filter (0.01-0.1 Hz) to reduce the effect of low frequency drift and high-frequency noise ^43^. Finally, the images were spatially smoothed using a Gaussian filter to decrease spatial noise (6 × 6 × 6 mm^3^ full width at half maximum).

### MVPA

#### ReHo and ALFF calculation

The ReHo and ALFF map of each subject was calculated with DPABI to evaluate local spontaneous activity and brain functional synchronization in the resting state. The ReHo map of each subject was generated by calculating Kendall’s coefficient of concordance between a single voxel and the 26 nearest neighbor voxels in a voxel-wise manner for the entire time series ^14^. Note that the ReHo calculation used rs-fMRI data that were not smoothed during preprocessing. The ALFF map of each subject was generated using the following steps. First, the imaging data were temporally bandpass filtered (0.01<f<0.08 Hz). Second, the time series of each voxel was transformed into the frequency domain to obtain the power spectrum, and the square root was calculated at each frequency of the power spectrum. Finally, the average square root was obtained across 0.01-0.08 Hz at each voxel and then taken as the ALFF value ^44^.

#### Kernel ridge regression (KRR)

Using the ReHo and ALFF maps, MVPA was implemented to predict IGD severity (i.e., IAT score) in the Pattern Recognition for Neuroimaging Toolbox (PRoNTo: http://www.mlnl.cs.ucl.ac.uk/pronto) ^45^. The KRR was utilized as the regression algorithm in the present study. KRR, which is a kernel-based approach, has very good generalization performance. The linear kernel method was used to map these implicit features into a high dimensional feature space ^46^. Then, ridge regression, a linear least square regression with Tikhonov regularization (regularization that penalizes the sum of squares of the weights) was applied to predict the IGD severity ^47, 48^. Here, the ReHo and ALFF maps were used as input data in the analyses. Each 3D image was transformed into a column vector of features, and each value corresponded to a single corresponding voxel intensity. Then, whole-brain models with features from images were constructed to investigate whether the ReHo/ALFF pattern could predict IGD severity.

#### Cross-validation method

To test the generalizability of the predictve models, leave-one-out cross-validation (LOOCV) was implemented. The method involved all subjects training the model, but one was left out to obtain an estimated model. The model was used to predict the behavior of the left-out subject. The above procedures were repeated n times (n= total number of subjects) to obtain a relatively unbiased estimate of generalizability. The performance of the established model was evaluated by two metrics, Pearson’s correlation coefficient (r) and the standard mean squared error (SMSE) ^49^. The correlation coefficient refers to the strength of a linear relationship between two variables. A higher correlation indicates better predictions. The SMSE refers to the mean of the squared differences between the predicted and true scores divided by the targets’ variance. The significance of these predictions was assessed with a permutation test of 1,000 permutations. That is, the same cross-validation procedure mentioned above was performed 1000 times with the label permuted across all the participants. The number of permutations that showed better performance (i.e., higher r or lower SMSE than the value obtained with the true target) was calculated. The p-value was computed by dividing the number by the total number of permutations (i.e., 1,000). The workflow for the MVPA is presented in Fig. 1.

**Figure 1.**
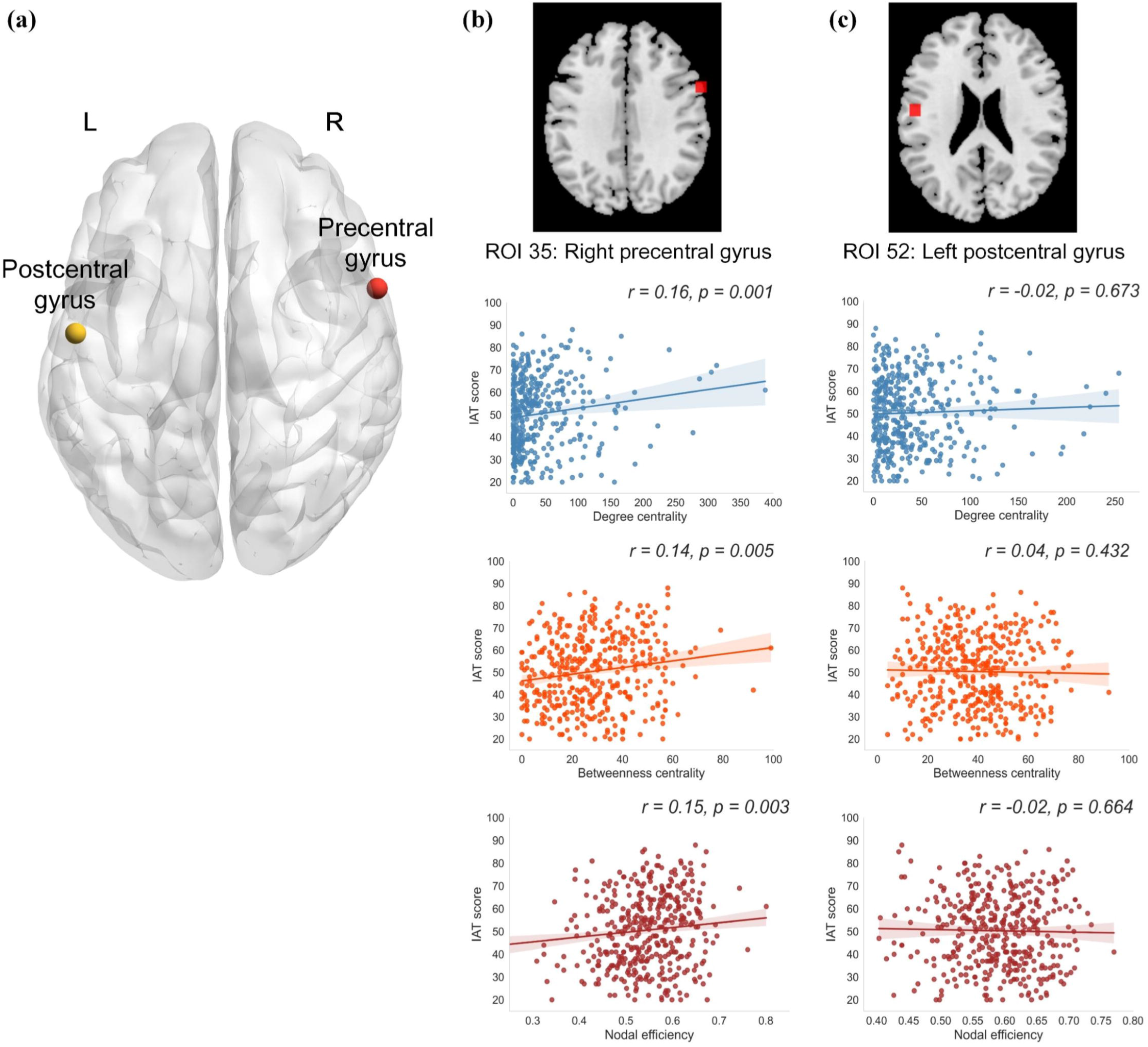
Workflow of multi-voxel pattern analysis. The ReHo and ALFF map were calculated based on rs-fMRI data and then were entered into predictive model as features to predict IGD severity. The KRR algorithm was applied to make prediction using cross-validated approach. The model performance was assessed by permutation test. IAT: Internet addiction test.

**Figure 2.**
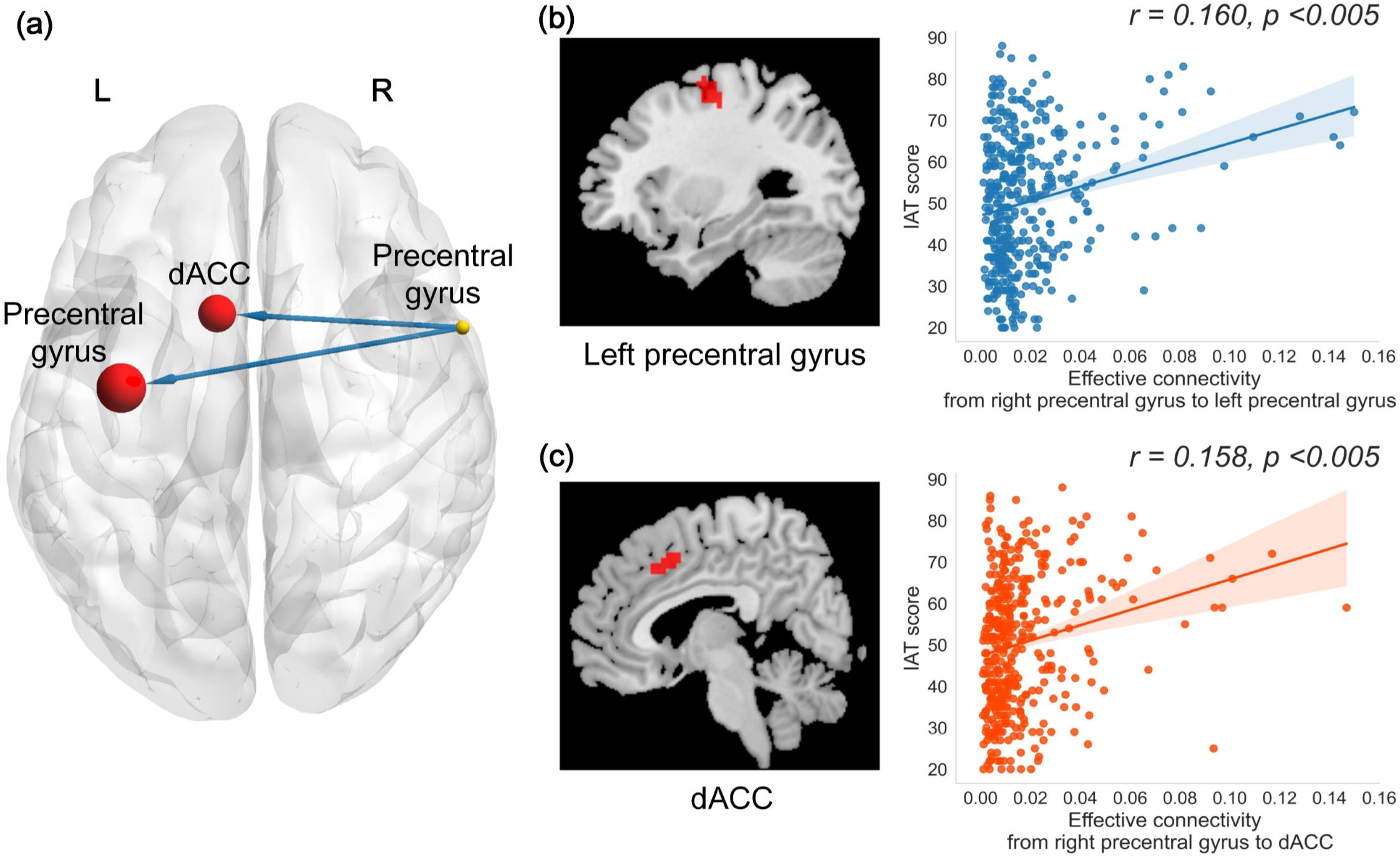
Performance of the model in predicting IAT score. Permutation distribution of the correlation coefficient (r) and standard mean squared error (SMSE) for the prediction analysis. The value obtained using the real scores are indicated by the red dash line. Higher *r* value and lower SMSE value indicate better performance of predictive models for ReHo and ALFF. \**p*<0.05.

### Region of interest (ROI) selection

A weighted image can be calculated by PRoNTo voxel-wise and ROI-wise for each model. To find the most weighted region in the whole brain, the Dosenbach 160-node atlas was applied to define the brain regions ^45, 50^. The ROI contributions to the predictive model were ranked in descending order, and regions that were 5% of the maximum (i.e., top 8) weighted were listed from predictive models that was based on the ReHo and ALFF patterns respectively. Regions that were presented in both top 8 lists were selected as ROIs in the current study.

### Graph theory measures

To test the significance of these ROIs in IGD severity across the brain network, graph theory was employed to characterize the topological properties of the ROIs. In graph theory, a topological brain network can be constructed by nodes and edges. Nodes refer to brain regions predefined by an atlas, and edges are defined as functional connectivity between two regions using Pearson’s correlation coefficient. Here, three nodal properties were computed to describe ROIs we selected in the predictive model. The degree centrality (DC) is the number of edges connected to a given node; it quantifies the information communication ability of nodes in the network ^51^. The betweenness centrality (BC) is the number of shortest paths passing through a given node and describes the effect of the node on the information transmission of other nodes ^52^. The nodal efficiency (NE) is the average inverse shortest path length between the given node and every other node, and it characterizes the efficiency of parallel information transfer by the node ^53^.

The nodes were defined using Dosenbach’s 160-node atlas, which was employed in the MVPA above. The edges were the functional connectivity between each pair of nodes, computed as Pearson’s correlation between the time courses of each pair of ROIs. First, the preprocessed data were entered to calculate correlation coefficients between each pair of ROIs, and subsequent correlation coefficients were normalized to Z-scores with Fisher’s r-to-z transformation. Thus, a weighted undirected functional connectivity matrix was generated, and it was converted to a graph network by considering a threshold T (set as 0.25 in the present study) to ensure that the stronger edges, in descending order, can enter the network construction ^54^. Then, the weighted network was binarized, so that DC, BC, and NE could be calculated for every node in the setting threshold. Finally, correlation analysis was conducted between these topological metrics of the ROIs and IGD severity. The procedures mentioned above were implemented in the GRaph thEoreTical Network Analysis toolbox (GRETNA: http://www.nitrc.org/projects/gretna/) ^55^.

### Granger Causality Analysis

Although GCA was originally developed in the field of economics to find the causal relationship between two time-courses, it has also been widely applied in neuroscience studies. Here, voxel-wise GCA was employed to evaluate effective connectivity related to selected ROIs. DynamicBC, a MATLAB toolbox, allows performance of GCA for rest-state fMRI data ^56^. The preprocessed 4D rest-state fMRI image was entered to calculate effective connectivity between ROIs and voxels within the whole brain. Thus, IN (i.e., information transmitted from other voxels to a given ROI) and OUT (i.e., information transmitted from a given ROI to other voxels) effective connectivity brain maps were generated separately for selected ROIs in each subject. Correlation analyses were conducted using IGD severity to determine how information flow through selected ROIs was influenced by IGD severity.

## Results

### MVPA results

The two predictive models (ReHo and ALFF) yielded similar and significant predictions of IGD severity (model based on ReHo: *r*=0.11, *p*(*r*)=0.030, SMSE=3.73, *p*(SMSE)=0.033; model based on ALFF: *r*=0.19, *p*(*r*)=0.002, SMSE=3.58, *p*(SMSE)=0.002).

For each model, the region weights were ranked in descending order. We found two regions, namely, the right precentral gyrus (60, 8, 34, x, y, z) and left postcentral gyrus (−54, 9, 23, x, y, z), were shown in the top 8 predictor lists of the two models and were therefore chosen as our ROIs.

### Graph theory analysis results

Graph theory analysis was applied to identify the important role of these highly weighted ROIs in IGD brain networks. Significant positive associations were found between IGD severity and all three graph theory metrics (i.e., DC, BC, and ND) of the right precentral gyrus (DC: *r* = 0.16, *p* = 0.001; BC: *r* = 0.14, *p* = 0.005; NE: *r* = 0.15, *p* = 0.003), whereas no significant result was found for the left postcentral gyrus (DC: *r* = 0.02, *p* = 0.673; BC: *r* = 0.04, *p* = 0.432; NE: *r* = 0.02, *p* = 0.664) (Fig. 3). These results indicated that the precentral gyrus may play a significant role in the whole-brain network. We reasoned that the precentral gyrus works as an information intermediary responsible for transmitting and integrating information in an IGD brain. To define these potential pathways, GCA was implemented using the precentral gyrus as our seed region.

**Figure 3.**
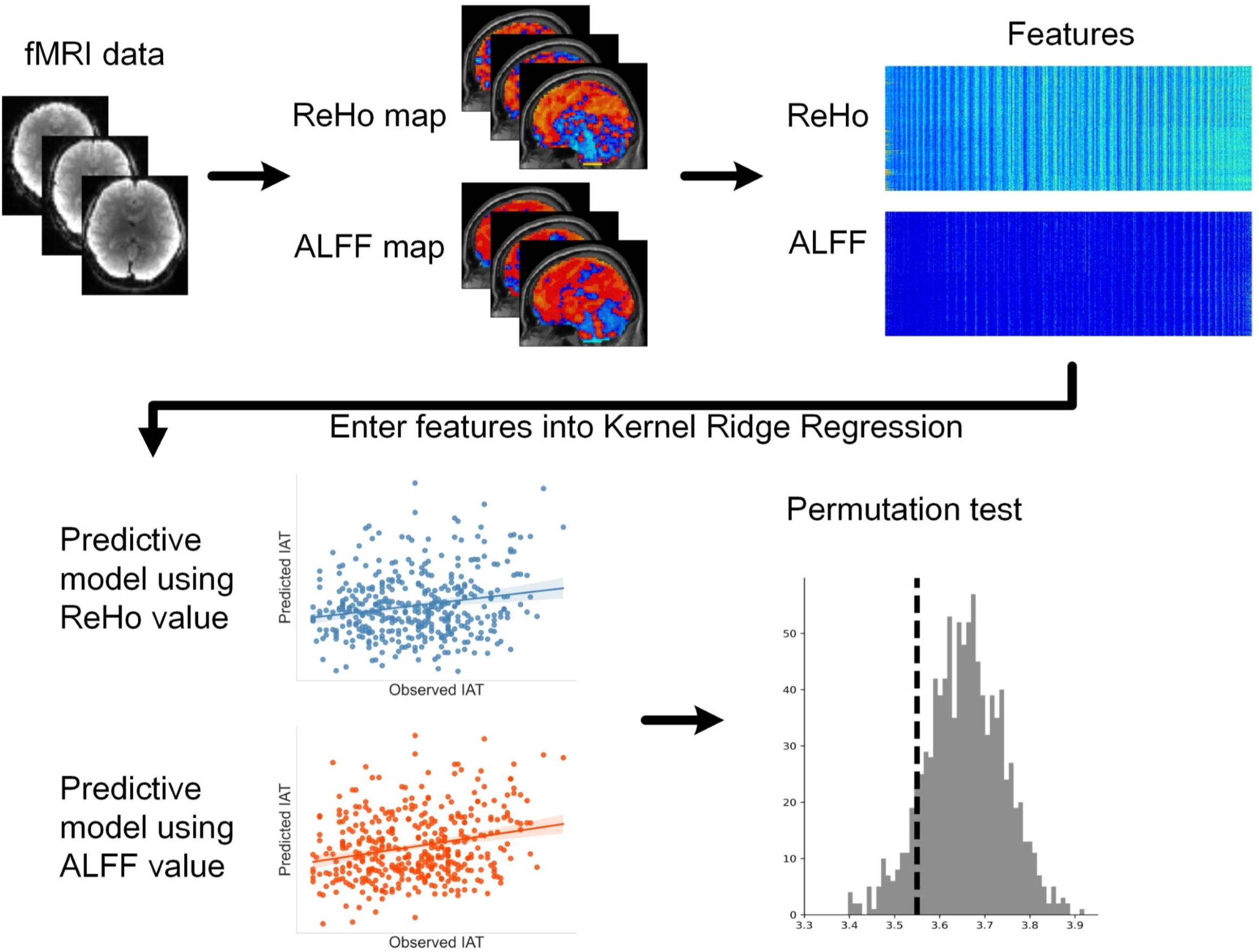
Two selected ROIs and results of graph theory analysis. The threshold *T* of network construction was set as 0.25. (a) Two highly weighted ROIs in both predictive models. (b) For the node of right precentral gyrus, the DC, BC, and NE are significantly related to IAT score. (c) For the node of left postcentral gyrus, the DC, BC, and NE are not associated with IAT score. IAT: Internet addiction test; DC: degree centrality; BC: betweenness centrality; NE: nodal efficiency.

### GCA results

Finally, GCA was applied to investigate the interactions between the right precentral gyrus and the whole brain. When correlating IGD severity (IAT score) with the effective connectivity results (input/output from right precentral gyrus), two effective connectivity that output from the right precentral gyrus to the left precentral gyrus (cluster size: 99 voxels; MNI coordinates: 33, −9, 57, x, y, z) and dACC (cluster size: 63 voxels; MNI coordinates: −9, 12, 45, x, y, z) were positively correlated with IAT score (*p* <0.005, GRF corrected) (Fig. 4).

**Figure 4.**
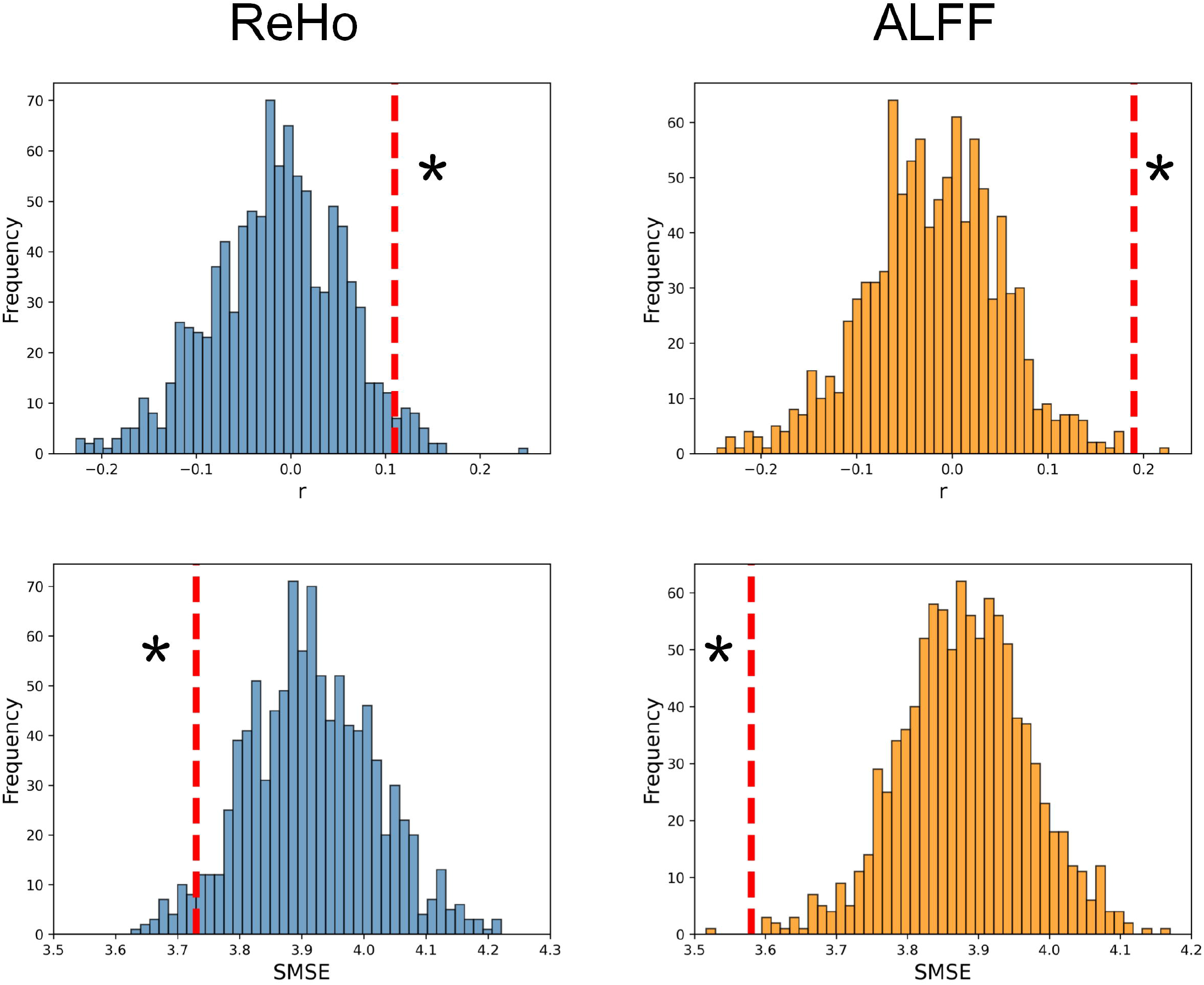
Result of Granger causality analysis. (a) Two effective connectivity were associated to IAT score. (b) Scatter plot shows the positive correlation between IAT score and the connectivity from right precentral gyrus to left precentral gyrus (r=0.160, p<0.005, GRF corrected). Each dot represents one subject. Semi-area represents 95% confidence interval for best-fit line. (c) Scatter plot shows the positive correlation between IAT score and the connectivity from right precentral gyrus to dACC (r = 0.158, p<0.005, GRF corrected). Each dot represents one subject. Semi-area represents 95% confidence interval for best-fit line. IAT: Internet addiction test; dACC: dorsal anterior cingulate cortex.

We reported the results using IAT score in selecting IGD, the results are of similar when using DSM-5 scores in selecting IGD. All results using DSM-5 scores were put into the supplementary materials.

## Discussion

In the present study, we combined machine learning techniques with rs-fMRI to identify distinct neural patterns in predicting IGD severity. The results showed that neural models represented by both ReHo and ALFF could significantly predict IGD severity at the individual level. In particular, the right precentral gyrus and left postcentral gyrus were highly weighted in both prediction models. Furthermore, graph theory analysis implied that the right precentral gyrus is an important node in the IGD brain network. Finally, GCA revealed that the effective connectivity between the right precentral gyrus and the left precentral gyrus and dACC was related to IGD severity.

### Neural patterns detected by MVPA can predict IGD severity

Previous studies have demonstrated that the resting-state neural patterns underlying IGD fit with its diverse behavioral characteristics. For example, subjects with IGD showed increased ReHo in brain regions involved in sensory-motor coordination ^18^. Furthermore, altered fALFF was observed in the cerebellum posterior lobe and superior temporal gyrus, which may be related to cognitive function and movement ^57^. Additionally, a recent meta-analysis found that abnormalities existed in several brain networks, including the default mode network, frontoparietal network, and attention network, in patients with IGD ^58^. Although these findings advance the understanding of the neural mechanism of IGD, the stability and reproducibility of these results still need to be evaluated with caution. The machine learning approach could decode complex brain patterns of mental disorders voxel-wise, which is more reliable than conventional fMRI analyses ^24, 59^. Here, we applied the MVPA method with two rs-fMRI metrics-ReHo and AlFF-to predict IGD severity. We found that both ReHo and ALFF can significantly predict addiction severity at the individual level. The results suggested that resting-state brain activities can be used to predict IGD severity effectively and imply the distinctive neural patterns underlying IGD. The brain activities in the informative brain regions of the predictive models were accompanied by changes in the severity of IGD, and these regions may play a key and distinct role in IGD. Moreover, consensus highly weighted brain regions were revealed by two predictive models, indicating that the results of MVPA are relatively robust and that neural signatures that can be detected by machine learning methods stably exist in IGD brains.

### Neural features in the precentral gyrus play a key role in predicting IGD severity

In both predictive models, the precentral gyrus and postcentral gyrus were reported as informative seeds for predicting IAT scores. The precentral gyrus and postcentral gyrus are the key regions of sensorimotor networks associated with integrating sensorimotor information and coordinating physical movement ^60, 61^. Atypical brain activities related to sensorimotor networks have been consistently indicated in IGD individuals previously ^20^. For example, IGD subjects showed enhanced ReHo values in brain regions associated with motor-sensory coordination ability and altered functional connectivity in sensory-motor related networks ^18, 33^. In line with these findings, the precentral gyrus and postcentral gyrus play a key role in motor functions, especially in coordinating hand movements, which is very important for internet game playing. The results of graph theory analysis further revealed the special role of the right precentral gyrus in IGD whole-brain networks. Elevated DC, BC, and NE of the right precentral gyrus were associated with greater IGD severity, indicating that the precentral gyrus may become a functional hub of the brain network that is responsible for information propagation, integration, and processing with increasing IGD severity ^62, 63^. Moreover, frequent computer game usage improves motor-visual coordination and perceptual-motor competencies ^64, 65^. Therefore, frequent internet game use, which is one of the features of IGD, may enhance interactions between the precentral gyrus and other regions to proficiently complete operations required by computer games^33^.

### Effective connectivity between the precentral gyrus and dACC related to IGD severity

Furthermore, we found that effective connectivity from the right precentral gyrus to the left precentral gyrus and the dACC was positively correlated with IGD severity. The effective connectivity from the right precentral gyrus to the left precentral gyrus may be related to the coordination of sensorimotor information between the left and right brain. Here, the increased effective connectivity between the left and the right precentral gyrus may indicate advanced game skills in individuals with IGD ^66^. Moreover, we found increased effective connectivity between the right precentral gyrus and the dACC, which is a node of salience networks in elevated IGD participants. The ACC is primarily associated with response selection ^67^, motivation ^68^, and reward assessment ^69^. The dorsal part of this region (i.e., the dACC) has been especially indicated, by an increasing body of research, to be dysfunctional among individuals with IGD ^70-72^. The dysfunction of the dACC may contribute to disadvantageous decision-making and cue-induced carving in addiction behaviors ^73, 74^. In the current study, we speculated that the pathway from the right precentral gyrus to the dACC may be responsible for transmitting and then integrating sensorimotor information into decision-making and reward evaluation. The enhancement of this pathway among individuals with IGD suggests more frequent communication between the right precentral gyrus and the dACC, leading to rapid reward assessment and risky decision-making ^75^. When subjects with IGD are exposed to game cues, they are strongly motivated to be involved in games and quickly respond to these cues, leading to their addiction and risky behavior. The effective connectivity results may reveal a neural circuit existing in the IGD brain. The enhancement of the circuit fits symptoms of IGD, i.e., that individuals are proficient in operating computer games that require extensive sensorimotor coordination and that they exhibit increased reward craving and strong motivation to play internet games ^7, 33^.

### Limitations

There were some limitations in this study. First, because this study was limited to a population of college students, most of whom are adults, it is difficult to determine whether the results would be the same in teenager samples. Future studies should examine the results in wider sample types. Second, we did not examine other highly weighted regions in the predictive model. Although some regions have a high weight only in a single model, they still have neurophysiological significance for IGD. How these regions’ neural activity is related to IGD severity needs full consideration and exploration.

## Conclusions

The current study demonstrated that resting-state neural activity could predict IGD severity at the individual level. As such, the findings may provide evidence to support the view that IGD has specific neural patterns and provide new insight into IGD. In addition, we discovered and verified the critical role of the precentral gyrus in IGD brain networks. The current study deepened our understanding of the neural mechanism of IGD and provides a potential target for physiological treatment interventions for IGD.

## ACKNOWLEDGMENTS

The current research was supported by the Zhejiang Natural Science Foundation (LY20C090005). The funding agencies did not have input into the writing of this manuscript.

## Authors’ contributions

Shuer Ye wrote the first draft of the manuscript. Shuer Ye and Ming Wang analyzed the data. Haohao Dong contributed to fMRI data collection. Guang-heng Dong designed this research. Guang-heng Dong and Qun Yang edited the manuscript. All authors contributed to and approved the final manuscript.

## Conflicts of interest

The authors declare that no competing interests exist.

## Notes

### Competing Interest Statement

The authors have declared no competing interest.

