## supplementary materials for "The neural features in the precentral gyrus predict the severity of internet game disorder: results from the multi-voxel pattern analyses"

**SUPPLEMENTARY MATERIAL**

**eResults.** MVPA and graph theory analysis results of DSM-5

**eDiscussion.** Discussion of study using DSM-5 score representing IGD severity.

**eTable1.** Participants demographics, and DSM-5 scores (N=365).

**eTable 2.** Top 8 predictors for IGD severity (DSM-5 score) and their relative weights in predictive power (percentage of the total weights) for predictive models based on ReHo and ALFF.

**eFigure 1.** Results for prediction of IGD severity (DSM-5 score).

**eFigure 2.** Two selected ROIs and results of graph theory analysis.

**eResults.** MVPA and graph theory analysis results of DSM-5

**MVPA results**

Two regression models (ReHo and ALFF) yield a similar and significant predictions on IGD severity (model using ReHo value: r=0.15, p(r)=0.004 SMSE=0.51, p(SMSE)=0.003; model using ALFF value: r=0.17, p(r)=0.009, SMSE=0.51, p(SMSE)=0.011, eFigure.1 ).

For each model, the region weights were ranked in descending order. We found two regions, namely left precentral gyrus (-38, -27, 60, x, y, z.) and right postcentral gyrus (-34, -39, 65, x, y, z), were shown in both top 8 predictors list of two models, thereby were chosen as our ROIs (eTable. 2).

**Graph theory analysis results**

Graph theory analysis was applied to identify the important role of these highly weighted ROIs in IGD brain networks. No significant correlation were found between IGD severity and all three graph theory metrics (i.e., DC, BC, and ND) of left precentral gyrus (DC: r =0.02, *p* = 0.761; BC: r = 0.03, *p* = 0.619; NE: r = 0.01, *p* = 0.887) and right postcentral gyrus (DC: r =-0.09, *p* = 0.099; BC: r = -0.06, *p* = 0.282; NE: r = -0.08, *p* = 0.119) (eFigure. 2). Due to no significant result was found between IGD severity and topological metrics of ROIs, we didn’t perform GCA in subsequent analysis.

**eDiscussion.**

We applied same analyses when using DSM-5 score to measure IGD severity and found both ReHo-based model and ALFF-based model can significantly predict DSM-5 score. These results consisted with the findings in study using IAT score to measure IGD severity. Also, precentral gyrus and postcentral gyrus were reported as consensus high-weighted regions, indicating IGD may associated with abnormality in sensory-motor related regions.

**eTable 1.** Participants demographics, and DSM-5 scores (N=365).

|  | Male  (N=203) | | | Female (N=162) | |
| --- | --- | --- | --- | --- | --- |
|  | Mean | SD | Mean | | SD |
| Age (years) | 21.58 | 2.63 | 20.98 | | 1.97 |
| Education (years) | 14.69 | 1.52 | 14.48 | | 1.09 |
| Gaming history (years) | 3.81 | 0.60 | 3.64 | | 0.67 |
| Gaming playing per weak(hours) | 6.98 | 3.30 | 7.03 | | 3.58 |
| DSM-5 score | 3.76 | 2.26 | 4.07 | | 2.05 |

SD: standard deviation; DSM-5: The Diagnostic and Statistical Manual of Mental Disorders, Fifth Edition.

**eTable 2.** Top 8 predictors for IGD severity (DSM-5 score) and their relative weights in predictive power (percentage of the total weights) for predictive models based on ReHo and ALFF.

|  | | Serial number ^a^ | | Anatomical region ^b^ | MNI coordinates | | | | | Weight (%) | Size(voxels) | Hemi |
| --- | --- | --- | --- | --- | --- | --- | --- | --- | --- | --- | --- | --- |
|  |  | |  |  | x | y | | z | |  |  |  |
|  | **ReHo model** | | 75 | Precentral gyrus | -38 | -27 | | 60 | | 2.42 | 19 | L |
|  |  | | 10 | Middle frontal lobe | -43 | 47 | | 2 | | 1.52 | 19 | L |
|  |  | | 50 | Superior frontal lobe | -26 | -8 | | 54 | | 1.40 | 19 | L |
|  |  | | 77 | Lateral occipital cortex | -24 | -30 | | 64 | | 1.39 | 19 | L |
|  |  | | 131 | Cerebellum | -34 | -67 | | -29 | | 1.21 | 19 | L |
|  |  | | 125 | Middle temporal lobe | -52 | -63 | | 15 | | 1.15 | 19 | L |
|  |  | | 68 | Superior temporal lobe | -54 | -22 | | 9 | | 1.14 | 19 | L |
|  |  | | 86 | Postcentral gyrus | 34 | -39 | | 65 | | 1.12 | 19 | R |
|  | **ALFF Model** | | 86 | Postcentral gyrus | 34 | | -39 | | 65 | 2.23 | 19 | R |
|  |  | | 107 | Inferior parietal lobe | 44 | | -52 | | 47 | 1.90 | 19 | R |
|  |  | | 87 | Middle cingulate cortex | 8 | | -40 | | 50 | 1.78 | 19 | R |
|  |  | | 75 | Precentral gyrus | -38 | | -27 | | 60 | 1.29 | 19 | L |
|  |  | | 35 | Precentral gyrus | 60 | | 8 | | 34 | 1.28 | 17 | R |
|  |  | | 7 | Anterior cingulate cortex | -6 | | 50 | | -1 | 1.27 | 19 | L |
|  |  | | 146 | Middle occipital lobe | -42 | | -76 | | 26 | 1.27 | 19 | L |
|  |  | | 17 | Superior frontal lobe | 23 | | 33 | | 47 | 1.25 | 19 | R |

a: The serial numbers are taken from Dosenbach 160-node atlas.

b: The anatomical regions were defined based on Automated Anatomical Labeling (AAL) atlas.

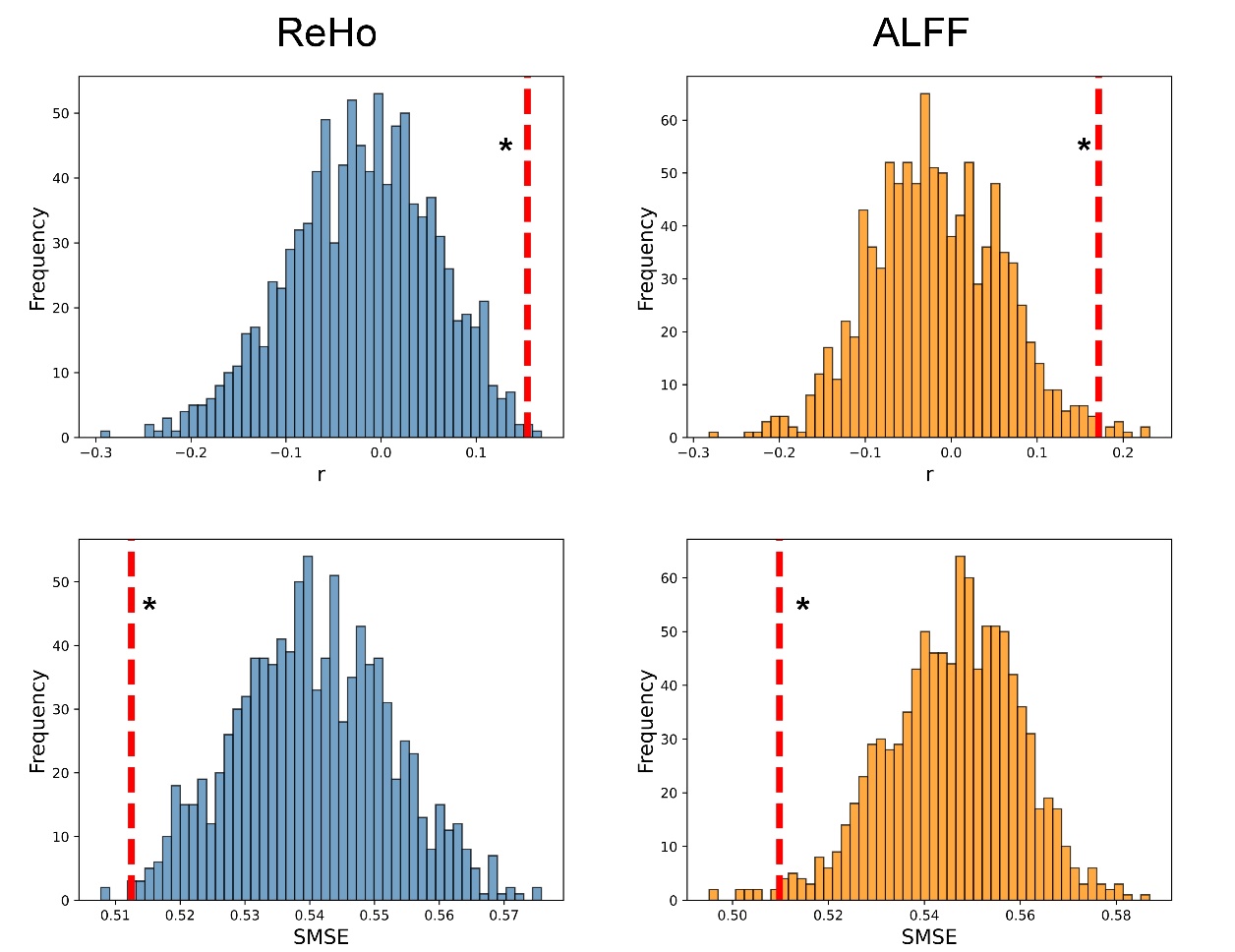

**eFigure 1. Performance of the** **model in predicting DSM-5 score.**

Permutation distribution of the correlation coefficient (r) and standard mean squared error (SMSE) for the prediction analysis. The value obtained using the real scores are indicated by the red dash line. Higher r value and lower SMSE value indicate better performance of predictive models for ReHo and ALFF. *p<0.05.

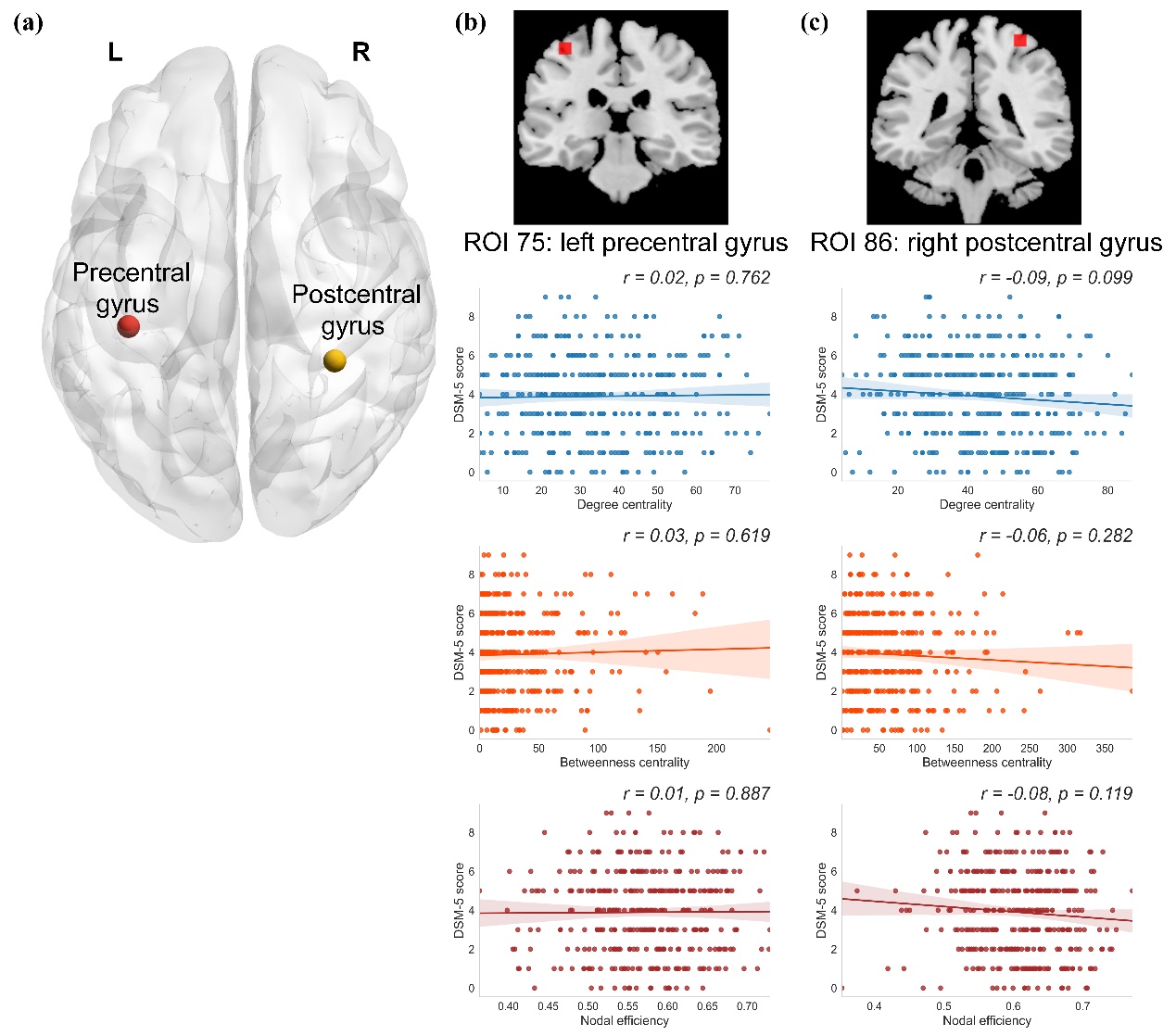

**eFigure. 2. Two selected ROIs and results of graph theory analysis.**

The threshold *T* of network construction was set as 0.25. (a) Two highly weighted ROIs in both regression models. (b) For the node of left precentral gyrus, the DC, BC, and NE are not associated with DSM-5 score. (c) For the node of right postcentral gyrus, the DC, BC, and NE are not associated with DSM-5 score. DC: degree centrality; BC: betweenness centrality; NE: nodal efficiency; DSM-5: The Diagnostic and Statistical Manual of Mental Disorders, Fifth Edition.
